# Flow transport and not ejection fraction determines left ventricular stasis in patients with impaired systolic function

**DOI:** 10.1101/2024.11.30.626199

**Authors:** Pablo Martinez-Legazpi, Javier Bermejo, Juan C. del Alamo

**Author notes:** **Corresponding Author:** Juan C. del Álamo, PhD., Department of Mechanical Engineering, Center for Cardiovascular Biology, University of Washington, Seattle, WA, US.

## Abstract

**Background:** Impaired left ventricular (LV) systolic function is a major risk factor for mural thrombosis and embolism, but LV ejection fraction (EF) poorly predicts these events, suggesting the existence of additional sources of variability. Advances in multi-dimensional flow imaging and patient-specific simulations have sparked the derivation of diverse metrics to assess blood stasis and transit efficiency. However, simple models to interpret these metrics and their dependence on chamber function are lacking.

**Methods:** We introduce queue models of LV blood transit connecting two common metrics of LV efficiency: flow component analysis and residence time (RT) mapping. These models yield closed-form expressions for the average RT of blood in the LV as a function of EF, direct flow (DF) —blood entering and leaving the LV in one cardiac cycle, and residual volume (RV) — blood persisting in the LV for >2 cycles. Models’ performance was tested against RT obtained from vector flow mapping in 332 subjects, including controls and patients with acute myocardial infarction (AMI), hypertrophic (HCM) and dilated (DCM) cardiomyopathy.

**Results:** Queue models revealed RT is increasingly sensitive to DF as EF decreases, contradicting the traditional view of large DF as a teleological advantage. Instead, RT is minimized when blood transits in a first-in-first-out (FIFO) manner, while DF short-circuits the FIFO pattern, prolonging RT for other flow components. FIFO models show a good performance to assess RT in the studied subjects, especially when accounting for patient-specific DF and RV, with R: 0.62 for the pooled data, 0.70 for control, and 0.60, 0.80 and 0.40 for AMI, DCM and HCM groups respectively.

**Conclusion:** By developing queue models of LV blood transit, and testing them on a large clinical database, we show large DF may contribute to increased blood stasis when EF is low. These models also explain why EF is a poor thrombosis risk marker in AMI and DCM.

## 1. Introduction

Increased blood stasis is a well-known risk factor for LV mural thrombosis. When combined with sources of endocardial damage and a systemic procoagulant states such as chronic heart failure (HF) or acute myocardial infarction (MI), intraventricular stasis places patients at a particular risk for intraventricular thrombosis and cardioembolism. Patients with non-ischemic dilated cardiomyopathy (NIDCM) have a 3 to 8 times higher ischemic stroke risk than age-matched individuals (1). Likewise, the age-adjusted risk of ischemic stroke increases by a factor of 30 in the month following an ST-segment elevation myocardial infarction (2). A low LV ejection fraction (EF) is thought to increase the risk of LV mural thrombosis because a reduced myocardial contractility may exacerbate intracavitary stasis (3). Consequently, EF inversely correlates with the incidence of stroke in HF patients and MI survivors (4, 5). However, this correlation excludes large demographic groups (5). Moreover, large clinical trials of anticoagulant agents on HF patients with reduced EF have produced contradictory results (6, 7). This evidence has led researchers to hypothesize that, in addition to global chamber systolic function, flow transport efficiency plays a determinant role on LV blood stasis (8).

Flow in the LV follows complex swirling patterns connecting the diastolic inflow and systolic outflow jets. The discovery of these patterns prompted numerous investigations focusing on their origin and teleology, in relation to ventricular function (9). In general, it is believed that LV flow patterns have a physiological function of washing out the chamber and preventing stasis (10). LV flow patterns define the pathlines of blood particles, partitioning the chamber into four regions with distinct transport dynamics: direct flow (DF), retained inflow (RI), delayed ejection (DE), and residual volume (RV)(11). These regions are heavily altered in cardiomyopathies, enlarging the regions of RV, with volumes of blood retained for more than two cycles (12).

DF, defined as the volume that enters and exits the chamber within the same cardiac cycle, has been traditionally viewed as an indicator of transport efficiency and it is affected by cardiomyopathies (12, 13). In NIDCM, increased LV chamber sphericity, together with changes in contractility result in larger swirling flow trajectories, reduced DF, and increased RV (14, 15). Although studied in less detail, changes in flow components in the setting of an AMI yield similar results (16, 17).

LV residence time (RT) is defined as the time spent by blood particles inside the chamber and can be used as a marker of blood stasis. In recent years, methods have been developed for mapping RT from vector flow data obtained either from echocardiography or from the phase contrast sequences of cardiac magnetic resonance (18). Although mean RT of blood in the LV and EF are inversely related (19), the former shows far better prognostic value for predicting mural thrombosis, silent brain infarcts, and cardioembolic strokes in the setting of NIDCM and AMI than the latter (20-22). RT is intrinsically related to the compartmentalization of blood transit through the LV chamber (23). Intuitively, increased RV and reduced DF components are associated with a high RT, but there is no mathematical representation of such correspondence (24, 25).

The present study was designed to assess the interplay between stasis, LV blood transit components, and LV global systolic chamber function. In particular, we aim to relate RT — as dependent variable, to DF and RV on one hand and EF on the other — as predictors. For this purpose, we developed simple queue models of LV transit. Based on these models, we argue that this relationship is less intuitive than previously thought. Significant model predictions show that LV RT becomes more sensitive to flow transport as EF decreases. We find that increasing the DF component results in higher RT for a constant EF, since it causes more incoming blood to exit the chamber immediately while retaining the resident blood pool for longer times. While simplistic, our models provide novel insights about the determinants of LV blood stasis and calls for caution when interpreting and comparing different metrics of LV flow transit. The clinical implications of potentially modifying intracardiac transport are highlighted.

## 2. Methods

### 2.1 Study Population

We prospectively selected 332 subjects from an existing database of previous studies (21, 22, 26, 27), including 90 patients with non-ischemic dilated cardiomyopathy (DCM), 93 patients with hypertrophic cardiomyopathy (HCM), 82 patients within the first 48h after acute myocardial infarction (AMI) and 67 healthy volunteers (Control). All subjects complied the following inclusion criteria: 1) age >18 y.o., 2) presence of sinus rhythm, 3) suitable apical ultrasonic window, and 4) absence of significant aortic regurgitation (grade *<* 2–3). Healthy controls were enrolled based on the absence of known or suspected cardiovascular disease, no diabetes or hypertension and normal electrocardiographic and ultrasound examinations. The Institutional Ethics Committee at Hospital Gregorio Marañón (Madrid, Spain) approved the studies, and all participants provided written informed consent.

### 2.2 Image Acquisition and Analysis

Ultrasound studies were performed using Vivid 7/E95 scanners and broadband transducers (GE Healthcare) following current recommendations (28). We obtained 2D sequences from parasternal and apical views to ensure full LV coverage to measure chamber volumes and ejection fraction. Pulsed-wave Doppler spectrograms were obtained at the level of the mitral tips and the LV outflow tract, and the temporal landmarks of each cardiac cycle were measured from spectral Doppler recordings using EchoPac software (v. 204, GE Healthcare).

### 2.3 Blood Transport & Stasis mapping

The two-dimensional (2D+t) blood flow field in the LV was obtained using vector flow mapping (VFM), as widely described elsewhere (29-31). Inputs for the VFM are a color-Doppler acquisition (∼10 beats) followed by a 2D cine-loop (∼5 beats) at high frame rate without moving the ultrasound probe. By integrating the fluid mechanics continuity equation, imposing no-penetration condition at the chamber walls, VFM yields the crossbeam flow velocity. The 2D+t blood flow fields from the VFM, 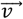, were used to integrate a forced advection equation to obtain the RT maps of blood inside the LV (18):

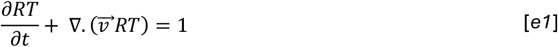

We integrated equation *e1* during 8 cycles, the estimate to washout the entire blood pool in a healthy LV (27), and collected the averaged value of RT at the end of last cycle (21, 22, 27). Equation *e1*, without the unity forcing, was also integrated both forward and backwards in time during two cycles, to track the advection of a scalar, ∅, allowing us to determine size and location of blood transport regions (DF, RI, DE and RV) at end-diastole (ED) (32, 33).

### 2.4 Queue Models for LV residence time

This section presents several models to estimate the mean RT in the LV, accounting for different LV transit patterns and microscopic mixing, using EF as the independent variable. The models are presented in order of increasing complexity, starting with the simplest mixing hypothesis and progressively incorporating flow information by adding the contributions of DF, given that *DF* + *DE* = *EF*, and RV to EF. Although some of these models obviously oversimplify blood transit in the LV, we believe that undergo the exercise of sequentially derive them helps to understand better how blood transits the LV. By comparing how different models fit different cohorts, we aim to shed light onto the dominant LV transit pattern in these groups.

#### 2.4.1 Perfect mixing transit model

The mean LV RT can be calculated as a function of the EF alone by assuming that blood inside the chamber achieves perfect microscopic mixing at the end of diastole. While this assumption does not accurately reflect flow transit inside the human LV (11), it allows to make simple model predictions that can serve as reference for the rest of the models developed in this study. It also provides a gentle introduction to the more complex transit models derived in this section. Following the diagram in **Figure 1A**, the mean LV residence time under perfect mixing can be obtained in closed form starting with the recursive formula

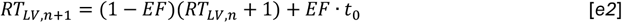

where the parameter *t*_0_ < 1 *cycle* represents the duration of diastole (more precisely, it is the mean RT of the blood that enters the LV each cycle at the end of diastole). This parameter also appears in all the other models and, while it may vary slightly from patient to patient, we fixed it to a constant value *t*_0_ = 0.7 cycles for the sake of simplicity. Following *e2* to the limit *n* → ∞, and noting that (1 − *EF*)^*n*^ → 0 in that limit, we obtain

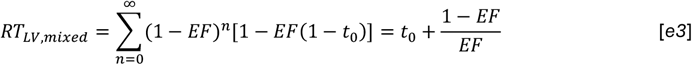

after some manipulation. When the EF approaches one, the model yields *RT*_*LV*_ ≈ *t*_0_. On the other hand, the model predicts that *RT*_*LV*_ becomes infinitely high in the limit when EF tends to zero.

**FIGURE 1:**
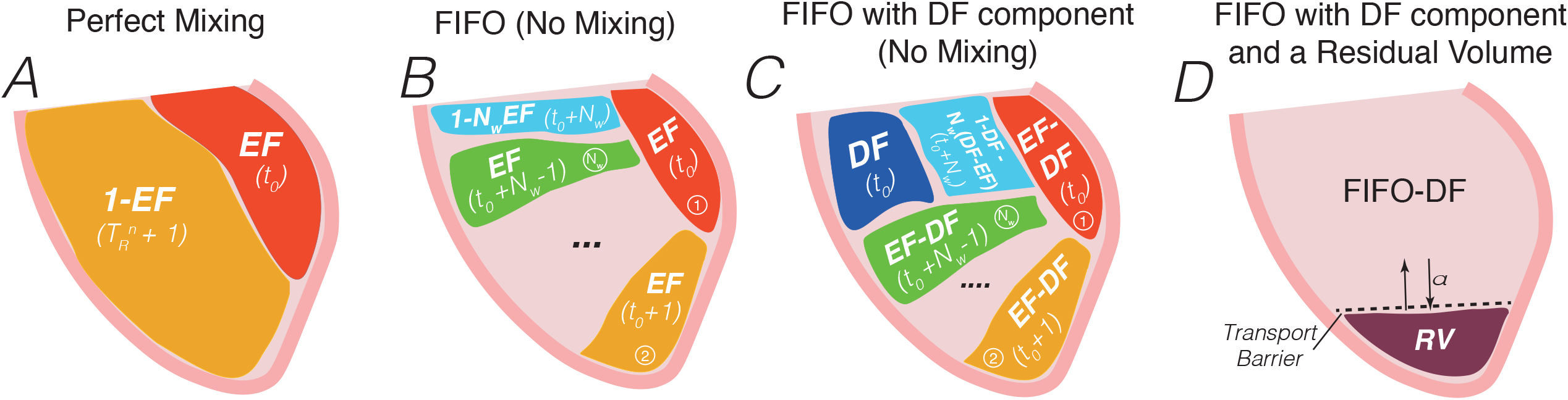
Sketch of the queue models. From left to right: **A-**Perfect Mixing, **B-**First-in-First Out (FIFO), **C-**FIFO with Direct Flow (FIFO-DF) and **D-**FIFO with DF and a residual volume (RV). The size of each region, scaled to the end-diastolic volume, is displayed in capital letters. The residence time for each region is shown in lowercase. EF: Ejection fraction, *N*_w_ : washout number, *t*_0_: diastole duration, *α*: fraction of blood within the residual volume that is exchanged with the rest of the left ventricle.

#### 2.4.2 Zero mixing with perfect FIFO transit model (FIFO)

For any given EF value, the mean RT in the LV is minimized when blood transits the chamber following a first-in first-out (FIFO) flow pattern that makes blood with the highest RT to be the first to exit the chamber. Qualitatively, the FIFO pattern is justified by the existence of diastolic vortices that redirect the inflow to outflow. The opposite transit pattern, i.e., last-in first out or LIFO, leads to infinite RT since it flushes incoming blood while indefinitely retaining blood already present in the chamber. For this reason, the zero-mixing LIFO model is trivial, and we will only consider the FIFO model here. Assuming zero mixing, the zero-mixing FIFO model partitions the LV chamber into 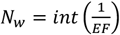 equal queued compartments of volume *V*_*i,EF*_ = *EF* · *EDV*, as shown in **Figure 1B**. We denote *N*_w_ the washout number as it measures the number of cardiac cycles required to clear the LV completely. RT inside each of the queued *N*_w_ compartments is *RT*_*i,EF*_ = *t*_0_ + *i* − 1 cycles. Since the *EDV* is usually not an exact multiple of *V*_*i,EF*_, there will be a smaller, residual ejection compartment with volume *V*_ΔE_ = *EDV* · (1 − *EF* · *N*_w_) and residence time *RT*_Δ_ = *t*_0_ + *N*_w_ (**Figure 1B**). The mean residence in the LV using this model, can be written as:

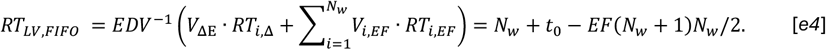

In *e4*, the only independent variable is EF and the parameter *t*_0_ is fixed, as described above. When *EF* = 1, *N*_w_ is also equal to one and we obtain *RT*_*LV*_ ≈ *t*_0_, like in the perfect mixing model. Below, we will see that the zero-mixing FIFO model is a particular case of a more general model that involves EF and DF when the direct flow component is fixed to be zero, i.e., *DF* = 0. On the other hand, the LIFO model is obtained when *DF* = *EF*.

#### 2.4.3 Imperfect FIFO transit with direct flow component (FIFO-DF)

We next consider a zero-mixing model where a fraction of the stroke volume is a direct flow compartment exhibiting LIFO transit. This imperfect FIFO model is characterized by two independent variables, i.e., *EF* and *DF*. Since *EF* = *DF* + *DE*, where the delayed ejection *DE* ≥ 0, the model is constrained to the *DF* ≤ *EF* half plane in the space of independent variables. As before *t*_0_ is considered constant. Following the same reasoning, we used in the ideal FIFO case one may obtain a closed-form relationship for the mean RT. However, in this case the washout number is 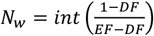, the equal queued compartments have volumes *V*_*i,EF,DF*_ = (*EF* − *DF*) · *EDV*, and the residual ejection compartment has a volume *V*_ΔE_ = *EDV* · (1 − *DF* − (*EF* − *DF*) · *N*_w_) (see **Figure 1C**). In this case, the LV mean residence time is given by

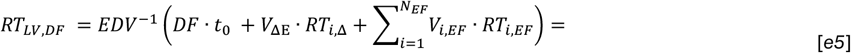

The rightmost side of this expression shows LV washout is delayed by the direct flow component and that the delay is roughly proportional to *DF* and inversely proportional to the square of the ejection fraction. It is also straightforward to see that *RT*_*LV,DF*_ becomes infinitely high when *DF* = *EF* because *N*_w_ → ∞. Also interesting, the model recovers that *N*_w_ = 1 when the ejection fraction becomes one, so that *RT*_*LV,DF*_(*EF* = 1) = *t*_0_ regardless of the direct flow component, converging with the other two models derived above.

#### 2.4.4 FIFO-DF with a transport barrier and residual volume (FIFO-DF-RV)

We expanded the FIFO-DF model described above to quantify how residual volumes isolated by transport barriers affect RT. Following the diagram of **Figure 1D**, the mean LV residence time in this case is given by

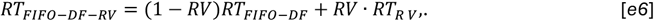

where *RV* is the residual volume component and *RT*_*RV*_ is the mean residence time inside the residual volume. Several working expressions can be obtained for *RT*_*RV*_, allowing for better interpreting this model and reducing its dependence on additional parameters. First, assuming that blood inside the RV has residence time high enough to be well mixed, the RT in the residual volume can be expressed as

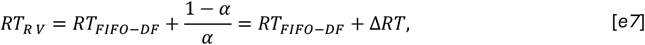

where *α* is the fraction of blood within the residual volume that is exchanged with the rest of the blood pool inside the LV. We note that this exchange’s contribution to *RT*_*FIF*0–*DF*_ was neglected for simplicity, since it only introduces a small, second-order effect. Another interpretation of the model is made possible by defining 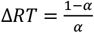 as the increment in RT inside the residual volume with respect to its value in the rest of the LV chamber. Combining the expressions above, we have:

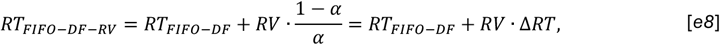

indicating that *RT*_*FIFO*–*DF*–*RV*_ = *RT*_*FIFO*–*DF*_ when there is no residual volume (*RV* = 0) or if *RV* is fully exchanged with the rest of the chamber every cardiac cycle (*α* = 1). On the other hand, *RT*_*FIFO*–*DF*–*RV*_ → ∞ when the residual volume is fully isolated and *α* → 0 (or equivalently, ∆*RT* → ∞). It is important to note that *RV* can be non-zero in the absence of transport barriers. For instance, a FIFO model with *EF* < 0.5 has a region with *RT*_Δ_ > 2 cycles, which constitutes a residual volume according to its original definition. However, this residual volume is in continuous transit rather than isolated beyond a barrier.

While the previous expression is useful to understand the model, it depends critically on an additional parameter (*α*) that is difficult to quantify. In an effort to minimize and clarify this dependence, we will estimate *RT*_*R V*_ under the assumption that it is determined by diffusion with an effective diffusion coefficient *κ* = *u*_*l*_, where *u*_*l*_ is the characteristic velocity of small-scale eddies in the LV and *l* is an effective mixing length (34, 35). Reasonable estimates for these quantities are *u*_*l*_ ≈ 0.1 *m*/*s* and *l* ≈ (*EDV* · *RV*)^1/3^ (36). We then take the simplified, 1D version of the governing equation for residence time with diffusion, under no flow conditions, and in steady state equilibrium (18),

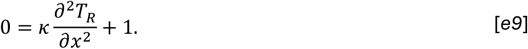

Equation *e9* can be solved imposing as boundary conditions ^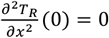^, i.e., zero diffusion of residence time and the LV wall, and *T*_*R*_(*l*) = *RT*_*FIFO*–*DF*_, i.e., that residence time at the interface of the residual volume matches the mean value in the rest of the chamber. The solution is *T*_*R*_(*x*) = *RT*_*FIFO*–*DF*_ − (*x*^2^ − *l*^2^)/(2*κ*). The spatially averaged value inside the residual volume is estimated as

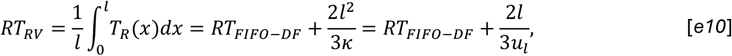

so that

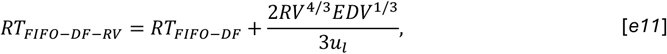

which is independent of the value of *α*. Similar to equation *e5*, this expression provides the LV washout delay associated to the residual volume.

### 2.5 Statistical analysis

Variables are described as mean and standard deviation unless otherwise specified. One-way analyses of variance (followed by Dunnett’s contrasts against the control group) were used to compare quantitative variables among the groups. Proportions and quantitative differences between 2 groups were assessed using χ2 and student t-tests, respectively. Statistical significance was established at the p< 0.05 level. Statistical analyses were performed using R version 4.3.

## 3. Results

### 3.1 Patient Characteristics

We analyzed a database of N=332 patients with different etiologies to evaluate the queue models presented in **Section 2.4**. Demographic and echocardiographic data are shown in **Table 1**. Compared to controls, the AMI and DCM groups were older and had lower ejection fractions (EF: 43 ± 9 % and 36 ± 11% vs. 63 ± 5%, p<0.001 for both). The HCM group had similar age as controls and their EF (65 ± 10%) was slightly higher. All patients had larger LV mass than controls (**Table 1**). The AMI and DCM groups had higher mean LV residence time than controls (RT: 3.1 ± 1.0 cycles and 2.2 ± 0.8 cycles vs. 1.7 ± 0.6 cycles, p<0.001) as well as larger residual volume (RV 52 ± 11% and 49 ± 15% vs. 32 ± 8%, p<0.001). AMI and DCM groups exhibited reduced direct flow (DF: 11 ± 7 % and 5 ± 5% vs 25 ± 8%, p<0.001). This reduction was especially drastic for DCM, for whom the mean DF decreased by a factor of 5 while the mean EF decreased by less than a factor of 2. HCM patients exhibited similar RT and RV as controls but had a larger DF component (DF: 35 ± 14% vs. 25 ± 8%, p<0.001), which was balanced by a lower retained inflow (RI: 15 ± 9% vs 20 ± %, p<0.001) and delayed ejection (DE: 19 ± 8% vs 23 ± 6%, p<0.001).

**Table 1.** Clinical & Imaging Data.

|  | Overall | Control | AMI | DCM | HCM | ANOVA-p |
| --- | --- | --- | --- | --- | --- | --- |
| N | 332 | 67 | 82 | 90 | 93 |  |
| Age (years old) | 55 ± 16 | 49 ± 18 | 60 ± 14* | 58 ± 15* | 52 ± 14 | <0.001 |
| Female Sex, n (%) | 126 (38%) | 35 (52%) | 15 (18%) | 37 (41%) | 39 (42%) | <0.001 |
| Ejection Fraction (%) | 50 ± 16 | 63 ± 5 | 43 ± 9* | 36 ± 11* | 65 ± 10* | <0.001 |
| End-Diastolic Volume (ml) | 105 ± 44 | 88 ± 25 | 100 ± 28 | 140 ± 53* | 84 ± 34 | <0.001 |
| End-Systolic Volume (ml) | 55 ± 39 | 33 ± 10 | 58 ± 21* | 92 ± 46* | 30 ± 19 | <0.001 |
| Stroke Volume (ml) | 54 ± 19 | 55 ± 16 | 49 ± 15 | 55 ± 18 | 57 ± 25 | 0.2 |
| E-wave velocity (cm/s) | 65 ± 21 | 70 ± 17 | 55 ± 17* | 62 ± 19* | 71 ± 25 | <0.001 |
| A-wave velocity (cm/s) | 65 ± 22 | 58 ± 18 | 69 ± 18* | 71 ± 22* | 62 ± 27 | <0.001 |
| LV mass (g) | 196 ± 82 | 121 ± 30 | 185 ± 44* | 197 ± 75* | 245 ± 97* | <0.001 |
| LV blood Residence Time (cycles) | 2.3 ± 1.0 | 1.7 ± 0.6 | 3.1 ± 1.0* | 2.2 ± 0.8* | 1.9 ± 1.3 | <0.001 |
| Direct Flow (%) | 19 ± 15 | 25 ± 8 | 11 ± 7* | 5 ± 5* | 35 ± 14* | <0.001 |
| Retained Inflow (%) | 19 ± 8 | 20 ± 7 | 17 ± 6 | 24 ± 8* | 15 ± 9* | <0.001 |
| Delayed Ejection (%) | 21 ± 7 | 23 ± 6 | 19 ± 6* | 22 ± 7 | 19 ± 8* | <0.001 |
| Residual Flow (%) | 42 ± 15 | 32 ± 8 | 52 ± 11* | 49 ± 15* | 31 ± 13 | <0.001 |

\*p<0.05 vs Control group in Dunnett's contrasts. LV: Left Ventricle. AMI: Acute Myocardial Infarction, DCM:
Dilated cardiomyopathy. HCM: Hypertrophic cardiomyopathy.

Figure 2. shows instantaneous LV flow streamlines, residence time maps, and flow component partitions from representative individuals of the Control, AMI, DCM, and HCM groups. It can be observed, in agreement with previous works, that DCM patients have larger and stronger swirling patterns than controls, while these patterns are weaker in HCM patients (26, 37). They also indicate that apical residual volumes associated with high RT are characteristic of LV flow patterns in HCM and especially AMI patients. In these isolated areas, mixing with the rest of the blood pool is limited and confined by a transport barrier. On the other hand, the residual volume associated with vortices seems to be in transit, as its residence time does not grow as much as in the one in apical residual regions. Taken together, these results suggest that flow in the LV of AMI, DCM, HCM patients, and controls follow distinct transit patterns in line with existing studies (11, 13, 23).

### 3.2 Queue Model Predictions

This section presents key queuing model predictions that shed light onto the relationship between mean LV residence time and parameters such as LV ejection fraction, direct flow component, and residual volume.

**FIGURE 2:**
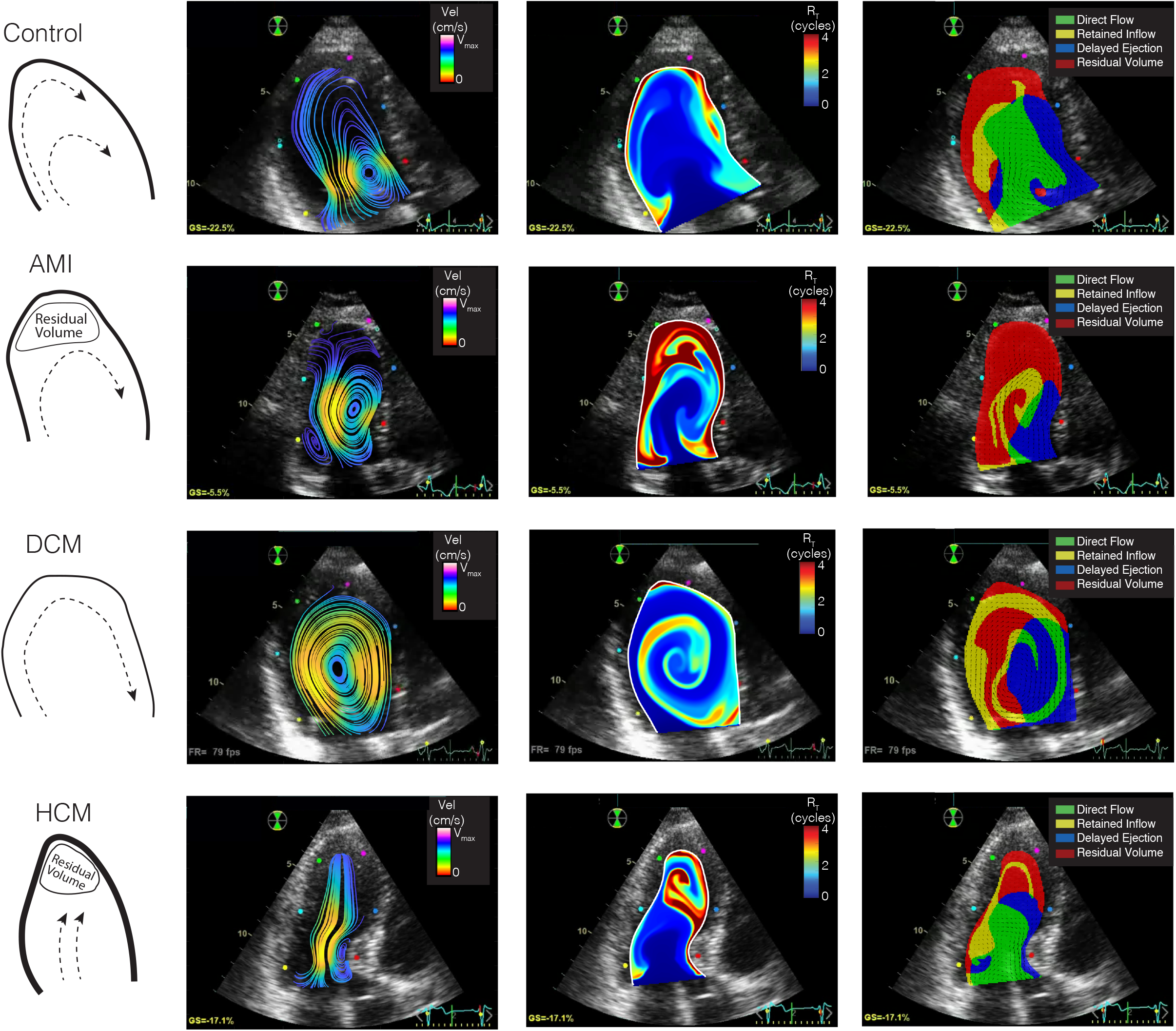
Examples of flow, RT and Transport. Sketch of main flow patterns (1^st^ column) and examples of the 2D+t flow fields (2^nd^ column), RT maps (3^rd^ column) and blood transport barriers (4^th^ column) for the control group (1^st^ row), AMI patients (2^nd^ row), DCM patients (3^rd^ row) and HCM patients (4^th^ row).

**Figure 3A** shows the dependence of RT on EF for the perfect mixing (black dashed line), zero-mixing FIFO (solid black line) and FIFO-DF (solid color lines) models. In the FIFO-DF case, the different curves are obtained by setting DF to a different constant value as indicated in the inset 2D map of RT vs. EF and DF. These curves are plotted in the range *DF* + 0.05 ≤ *EF* ≤ 1 to allow the delayed ejection *DE* = *EF* − *DF* to be at least 5% of the end-diastolic volume. In all curves, the mean RT increases as EF decreases consistent with our expectations. However, this dependency has interesting, previously unobserved features that emerge in this model.

**FIGURE 3:**
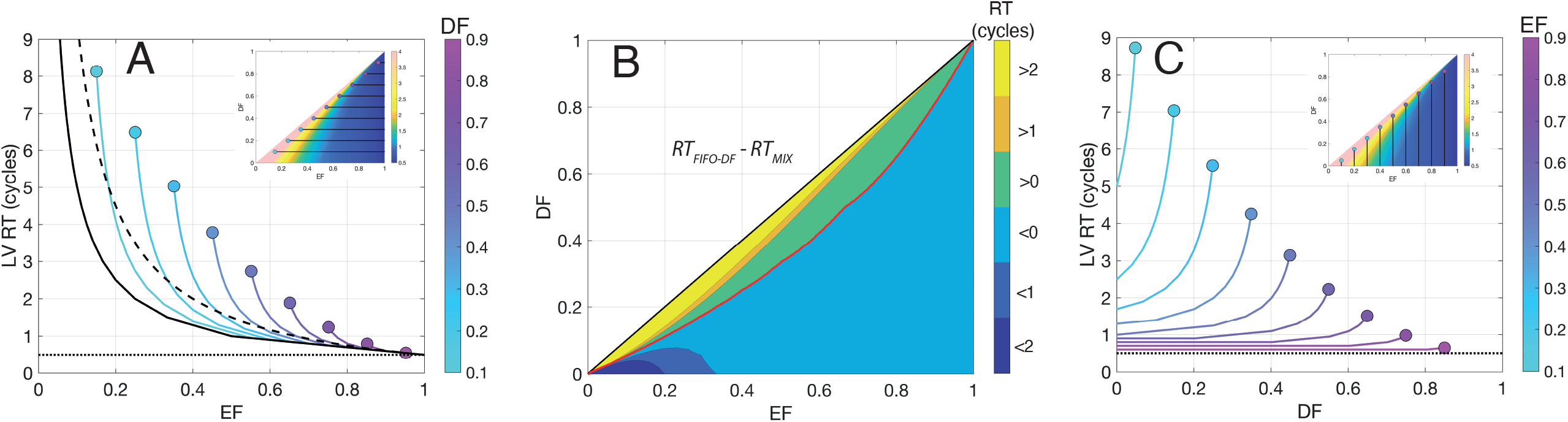
Model prediction. Panel A: Evolution of the RT as a function of EF for different values of DF in the FIFO-DF model. Solid line: zero-mixing FIFO model. Dashed-Line: Perfect mixing model. Dotted line: *t*_0_. **Panel B**: Subspace of the difference of RT obtained by FIFO-DF model and the perfect mixing model as a function of EF and DF. **Panel C**: Evolution of RT as a function of DF for different values of EF. Dotted line: *t*_0_.

The first important feature is that, as *EF* approaches 1, all the curves converge to the same point *RT*(*EF* = 1) = *t*_0_. This result implies that the mean RT is relatively insensitive to flow patterns and is almost uniquely determined by ejection fraction in ventricles with good contractile function. In contrast, the curves diverge significantly as *EF* decreases, suggesting that LV flow patterns are a critical determinant of blood stasis and *EF* is not sufficient to quantify RT in weakly contracting ventricles. A second interesting feature is that the FIFO transit pattern obtained for *DF* = 0 yields lower residence time than the perfect mixing model for all *EF* values. However, the FIFO-DF curves intersect the perfect mixing curve when DF>0, producing longer residence times than perfect mixing as *EF* decreases below the intersection point. This prediction suggests that mixing the incoming and resident blood pools may prevent stasis in LVs that contract poorly or have a large direct flow component. To analyze this possibility in more detail, **Figure 3B** maps the difference between the residence times obtained with FIFO-DF and perfect mixing transit, RT_*FIFO-DF*_ *- RT*_*mix*_, versus EF and DF. This figure shows that mixing becomes more effective at clearing the LV than FIFO-DF when the direct flow component increases beyond the value *DF* ≈ *EF*(1 + *EF*^2^)/2. Moreover, it shows that the difference in residence times can be significant for low EF values.

The queue model predictions presented so far suggest that increasing the direct flow component for a constant EF may worsen transit efficiency and increase LV stasis. This idea is tested in **Figure 3C** by plotting mean RT curves versus DF for different EF values, obtained as indicated in the inset *TR(EF,DF)* map. Similar to **Figure 3A**, we considered the range 0 ≤ *DF* ≤ *EF* + 0.05 to allow the delayed ejection to be at least 5% of the end-diastolic volume. For high EF values, RT is almost independent of DF, and differs little from the minimum value *t*_0_, confirming that the LVs with high contractile function are washed out efficiently regardless of flow patterns. On the other hand, the curves corresponding to low EF values grow with DF and become steeper as DF increases and EF decreases. These data support that LV blood stasis increases with the direct flow component for constant EF, especially as DF approaches EF. In this theoretical limit, there is no delayed ejection or retained inflow, the transit pattern becomes LIFO, and the LV entire blood pool that does not transit the LV within a cardiac cycle remains indefinitely inside the chamber.

### 3.3 Evaluation of Queue Models Using Clinical Data

We tested whether the key predictions of LV transit queue models reported above are reproduced in a clinical database of 332 individuals. **Figure 4A** shows the probability distributions of RT conditional to EF from all study subjects aggregated in three groups depending on the patient-specific value of direct flow: low DF (DF<0.1, top panel), intermediate DF (0.1<DF<0.3, middle panel), and high DF (DF>0.3, bottom panel). The dependence of these distributions on EF is reasonably well captured by the FIFO-DF queue model when DF is set to each group’s median value (i.e., 0.05, 0.2, and 0.4 for the low-, intermediate-, and high-DF groups). Patient data confirms that RT increases as EF decreases and this decrease is less pronounced in patients with a lower DF component. To focus on the DF dependence of residence time, we plotted the probability distributions of RT conditional to DF for our study population (**Figure 4B**). In this case, the data was also aggregated in three groups, depending on the patient-specific value of ejection fraction: low EF (EF<0.1, top panel), intermediate EF (0.3<DF<0.45, middle panel), and normal EF (EF>0.45, bottom panel). As in **Figure 4A**, the FIFO-DF model predictions for each group’s median value of EF are included for reference. Once again, the patient distribution followed well the FIFO-DF model predictions. In particular, the steep increase of RT with DF for low ejection fraction was distinctly observed in the clinical data. Additionally, a clear decline in the sensitivity of RT to DF was adequately identified in the patient measurements.

**FIGURE 4:**
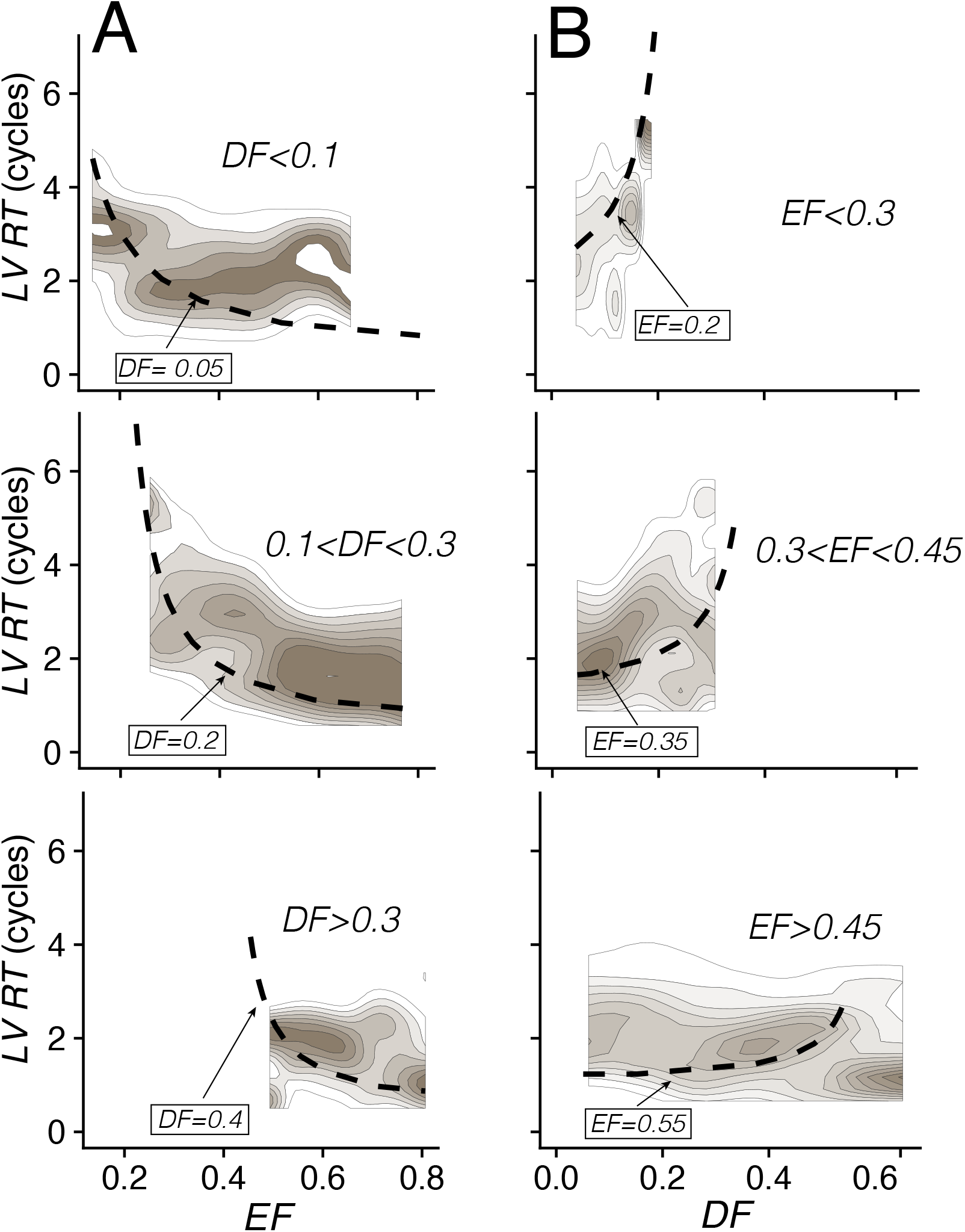
Residence time in the studied subjects. Panel A: Hexagonal heatmap plot, shaded by data density, of RT in all the subjects as a function of EF. Dashed-Line represents the perfect mixing model. Solid line is the averaged patient-specific FIFO-DF prediction, and the dotted line is the averaged patient-specific FIFO-DF-RV prediction. **Panel B**: Conditional probability density plot of RT in all the subjects as a function of EF. Data is split for low (<0.1), moderate ([0.1, 0.3]) and high (>0.3) DF. **Panel C**: Conditional probability density plot of RT in all the subjects as a function of DF. Data is split for low (<0.3), moderate ([0.3, 0.45]) and high (>0.45) EF. In both panels, the dashed line represents the FIFO-DF model for the median of the data shown.

Finally, we compared the mean RT predicted by the queue models using each patient’s EF, DF, and RV values obtained by echocardiography, with the patient’s RT obtained by echocardiographic VFM (**Figure 5**). The goal of this comparison was to test two hypotheses. First, that LV blood transit in DCM patients approximately follows a FIFO pattern where residual volume (blood lingering for more than two cycles inside the LV) is in transit. Second, that HCM and AMI patients have residual volume excluded from the main flow path that explains some of these patients’ elevated RT despite their normal or moderately impaired EF. **Figure 5A** displays the measured residence times versus the values predicted with the perfect mixing model that only depends on EF. This model offered a modest compromise fit for the entire population, yielding a correlation coefficient of *R=0*.*37* and a root mean-squared error *RMSE=*1.07 cycles. Moreover, it did not represent any group particularly well, overestimating the RT of DCM patients and underestimating the RT of the other three groups (**Table 2**). FIFO and FIFO-DF models were most effective at reproducing the residence time, especially in the DCM group (*R=0.61* and *0.64, RMSE=0.68* and *0.62 cycles*, respectively), although they severely underestimated the RT of the HCM and AMI groups (see **Figure 5B G C** and **Table 2**). The FIFO-DF-RV model shown in **Figure 5D** predicted the RT of the AMI and HCM groups significantly better than the other models (*R=0.60* and *0.40, RMSE=0.78* and *0.82 cycles*). However, it overestimated the residence time of the DCM group considerably (*RMSE=1.31 cycles)*. Overall, these results confirmed our hypotheses about the impact LV transit patterns have in the RT in the different cardiac conditions. This latter model was found also to be the most representative of the control group (*R=0.70, RMSE=0.43 cycles*).

**Table 2.** Model Performance.

| **Model** | **Pooled Data** |  | **Control** |  | **AMI** |  | **DCM** |  | **HCM** |  |
| --- | --- | --- | --- | --- | --- | --- | --- | --- | --- | --- |
|  | *R* | *RMSE* | *R* | *RMSE* | *R* | *RMSE* | *R* | *RMSE* | *R* | *RMSE* |
| Perfect Mixing | 0.37 | 1.07 | 0.10 | 0.72 | 0.28 | 1.36 | 0.63 | 1.11 | 0.10 | 0.77 |
| FIFO | 0.20 | 1.72 | 0.12 | 0.97 | 0.10 | 2.22 | 0.61 | 0.68 | 0.18 | 2.33 |
| FIFO-DF | 0.37 | 1.15 | 0.09 | 0.84 | 0.26 | 1.72 | 0.64 | 0.62 | 0.09 | 0.86 |
| FIFO-DF-RV | 0.62 | 0.88 | 0.70 | 0.43 | 0.60 | 0.78 | 0.80 | 1.31 | 0.40 | 0.82 |

R: Correlation Coefficient. RMSE: Root Mean Squared Error. AMI: Acute Myocardial Infarction, DCM: Dilated cardiomyopathy. HCM: Hypertrophic cardiomyopathy.

**FIGURE 5:**
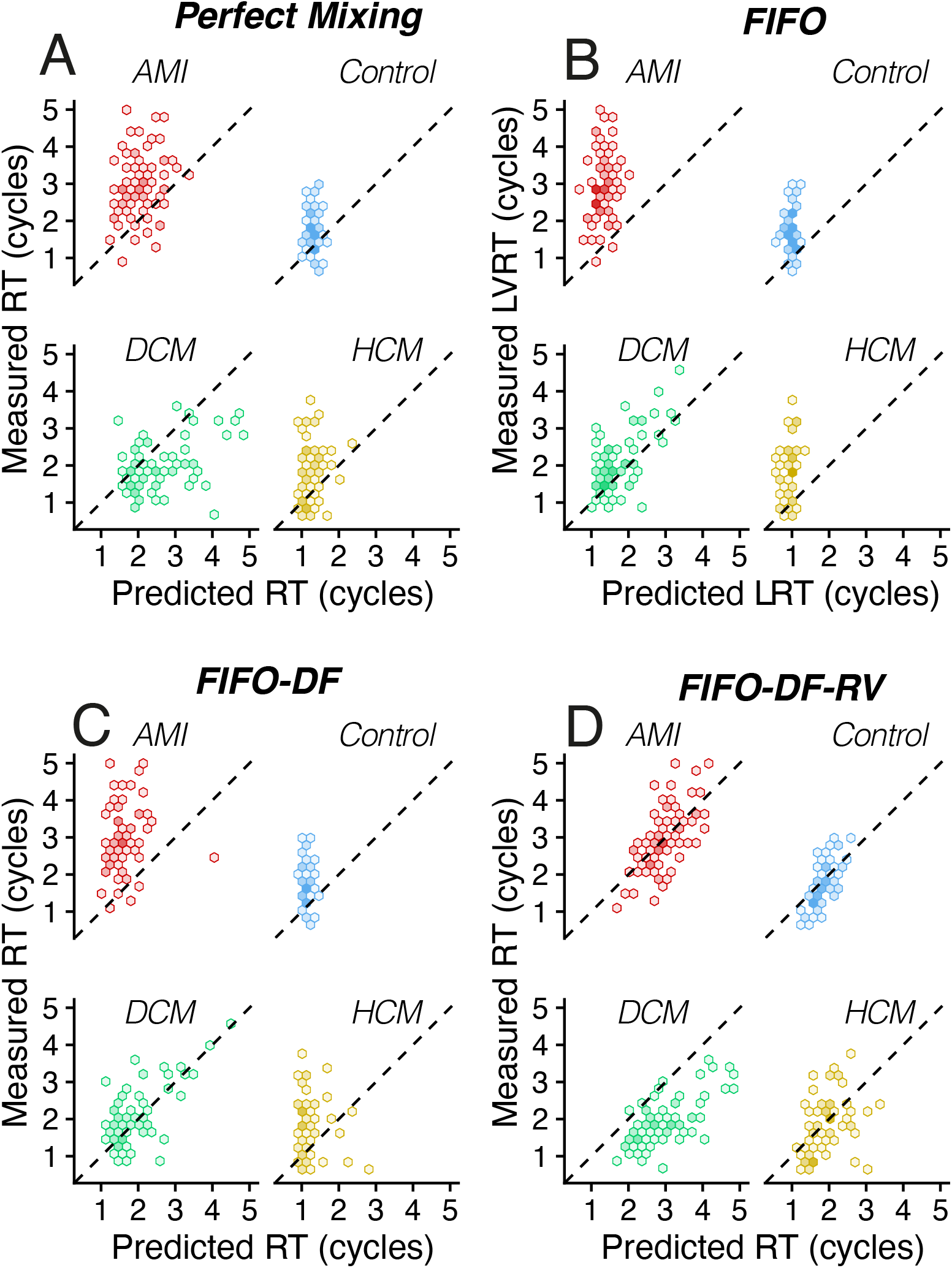
Hexagonal heatmap plot, shaded by data density, of the prediction and measured RT for the Perfect Mixing (**Panel A**), zero-mixing FIFO (**Panel B**), FIFO-DF (**Panel C**) and the FIFO-DF-RV (**Panel D**) models respectively. Data is colored by group. AMI: Acute Myocardial Infarction, DCM: Dilated cardiomyopathy. HCM: Hypertrophic cardiomyopathy.

## 4 Discussion

The LV harbors complex flow patterns hypothesized to organize short pathlines connecting the diastolic inflow jet with the systolic outflow (9). In diseased hearts, the impairment of systolic chamber function and the disruption of the flow path lead to increased blood stasis and thrombosis (38-40). Since EF is not independently predictive of LV thrombus or stroke (5, 21, 22, 41), a growing interest in clinical flow imaging and LV transit analysis has surged in the past decade. This trend has led to numerous analyses of intracardiac flow, using 1D and 2D fields from ultrasound data and 3D fields obtained either through the phase contrast sequences of the MRI or, more recently, to computational models using patient-specific anatomies (8, 12, 16, 26, 42, 43). In contrast with the rapidly growing body of high-fidelity clinical and simulation data, there are few theoretical models to help interpret such complex data. One-dimensional models of the LV inflow jet have produced metrics like the vortex formation time or the E-wave propagation index, which have been associated with LV disease state and mural thrombosis risk (44, 45). However, we lack models that consider the complete path of blood from LV inflow to LV outflow. Ideally, such models should incorporate global chamber function and allow for relating blood stasis with transit metrics derived from multi- dimensional imaging or simulation modalities.

In this work, we present queue models of LV transit to examine the relationship between residence time, LV ejection fraction, and the direct flow (DF) and residual volume (RV) components (11). We found this relationship less intuitive than previously thought for several reasons. First, flow transport efficiency affects residence time differently depending on global systolic chamber function. It is evident that a ventricle with an EF near 100% cannot have extensive stasis regardless of intraventricular flow patterns because most of its volume is cleared each cardiac cycle. Also, in LVs with reduced EF, stasis can still be moderate if the LV flow establishes a continuous path for all incoming blood to reach the LV outflow tract within a few cardiac cycles. Conversely, stasis can be large for normal EF values if a transport barrier excludes significant regions from the main transit path. Along these lines, the definition of RV as blood with RT> 2 cycles may cause ambiguity between blood in slow transit and truly stagnant blood (46). Besides, direct flow cannot be understood without considering systolic function since *EF* = *DF* + *DE* ≥ *DF*. This interdependency may obscure the interpretation of flow components in healthy and diseased groups. The low values of DF reported in diseased ventricles (8, 11, 13, 16, 17, 24, 32, 33, 47) could merely mirror their lower EF instead of indicating changes in flow efficiency. Likewise, a concurrent increase of DF and EF is observed following dobutamine infusion (47).

Consistent with the ideas discussed above, our queue models show that RT does not only increase as EF decreases. It also becomes sensitive to additional parameters that reflect the overall intraventricular flow pattern and flow-induced mixing. Notably, these models suggest the LV mean residence time increases with DF for a constant value of EF. While perhaps not intuitive, this finding reflects that increasing direct flow promotes more incoming blood to exit the chamber immediately while forcing the resident blood pool to be retained longer. The trend is also observed in clinical data when plotting RT versus DF for constant EF values.

The queue models of this work offer mathematical expressions for RT that can be particularized to patient-specific ejection fraction, direct flow, and residual volume values. While the models’ crudeness and involved parameterization compromise their predictive power, comparing their results with measurements can be informative. For instance, we found that including a transport barrier was essential to reproduce patient-specific RT values in the AMI group. In line with this finding, significant stagnant regions are often found near the LV dyskinetic apex in AMI patients (48). In contrast, low-DF FIFO models performed best in the DCM group, consistent with the large swirling pattern typically found in enlarged ventricles dominating blood transit. In dilated ventricles with low EF, this large-scale swirl allows for the entire blood pool to slowly transit from inflow to outflow, even if some of this blood is labeled as residual volume, using the definition RT > 2 cycles. Previous works highlighted the relevance of this flow structure, proposing it conducts mechanical energy efficiency (10, 45, 49) in connection with the reduction of the DF component (50). On the other hand, since the FIFO transit pattern achieves minimum residence time at any given EF, our models and data suggest that, under particular geometries, ventricular dilation may favor a blood flow pattern protective against mural thrombosis. This idea is supported by patients with DCM having lower LV RT than AMI patients when matched by EF. Whether this phenomenon reflects functional or evolutionary adaptations that favor blood clearance over mechanical energy conservation is debatable, but it underscores that not all forms of remodeling in cardiac disease are equally detrimental.

### 4.1 Clinical Implications

The models and data presented here explain why EF alone is a poor imaging biomarker to select patients for primary prevention of LV mural thrombosis in patients with impaired systolic function. Thus, it is likely that suboptimal criteria for patient selection may have hampered a favorable risk-benefit ratio of landmark clinical trials of preventing cardioembolic risk in patients with ischemic and nonischemic dilated cardiomyopathy (51, 52). Dependency on functional barriers that modulate intraventricular flow transit are key determinants of the risk of mural thrombosis in patients with abnormal LVs. LV regional wall motion, atrioventricular electrical coupling, and LV inflow geometry are well-known determinants of intraventricular flow transit. Consequently, our results explain why therapeutic procedures such as cardiac resynchronization therapy (32), mitral edge-to-edge repair (53, 54) and mitral valve replacement (55) may induce intraventricular stasis despite a favorable effect on LV EF. For the reasons discussed above, this will not be an issue in patients with normal EF, even in the presence of overt regional wall motion abnormalities (22). However, in patients with a low EF, flow imaging modalities are necessary to measure flow transit and/or stasis and, consequently, to account for the flow-mediated factors that trigger intraventricular thrombosis.

### 4.2 Limitations

All the FIFO models presented in this manuscript incorporate a free parameter, *t*_0_, which roughly represents the duration of diastole normalized with the cardiac period. The models are not overly sensitive to *t*_0_, since this parameter is an additive constant that sets the minimum value of RT. In principle, this parameter could be adjusted to fit our echo RT measurements, but we set it to a constant value for simplicity. Along the same lines, the FIFO-RV model introduces an additional free parameter *α* that accounts for the fraction of blood within the RV that is exchanged each cardiac cycle. To eliminate the model’s dependence on this parameter, we derived a simple phenomenological extension that depends on end-diastolic volume, residual volume, and a characteristic velocity, *u*_*l*_. This parameter can have a more significant effect RT than *t*_0_ because it sets the slope of the residual volume contribution (see *e11*). For simplicity, we set *u*_*l*_ equal to a constant value taken from the existing literature across all patients.

The limitations of the VFM modality, used to evaluate our models with clinical data, have been discussed thoroughly in previous publications (29, 56-58). In short, VFM assumes planar flow in the long-axis apical view of the LV, an approximation that has been justified by several groups (29, 56-58). Also, flow compartments and residence time have been reported to be well captured using VFM when compared to the reference 4D flow MRI approach (33).

Finally, in the queuing models, RT was calculated assuming mixing and transport processes span during infinite number of cycles *N*_*cycles*_ → ∞. However, the number of cycles used to compute RT from VFM data is necessarily finite. We selected *N*_*cycles*_ = 8 based on our observation that RT remains unchanged in normal LVs after 8 cycles (27). We ensured that this discrepancy did not cause significant differences by also evaluating our queue models for *N*_*cycles*_ = 8 (Supplemental Figure 1). Except for unrealistically low EF values, the models produced similar results for *N*_*cycles*_ = 8 and *N*_*cycles*_ → ∞.

## 5. Conclusions

We present simple queue models of LV transit to study how chamber function and flow patterns affect blood stasis. Intracardiac stasis becomes increasingly sensitive to flow patterns as EF decreases, which explains why EF is a poor predictor of LV thrombus and stroke in patients with impaired systolic function. Our models suggest that LV transit efficiency is paradoxically impaired when more incoming blood exits the chamber immediately, thus retaining the pool of resident blood longer. Particular flow patterns—like swirling flows in dilated spherical ventricles— are related to lower values of RT and, consequently, with lower cardioembolic risk.

## Notes

### Competing Interest Statement

The authors have declared no competing interest.

