## Supplementary Materials for "Flow transport and not ejection fraction determines left ventricular stasis in patients with impaired systolic function"

### The role of flow transport and chamber function on left ventricular stasis

#### Supplemental Material

#### Supplemental Figures

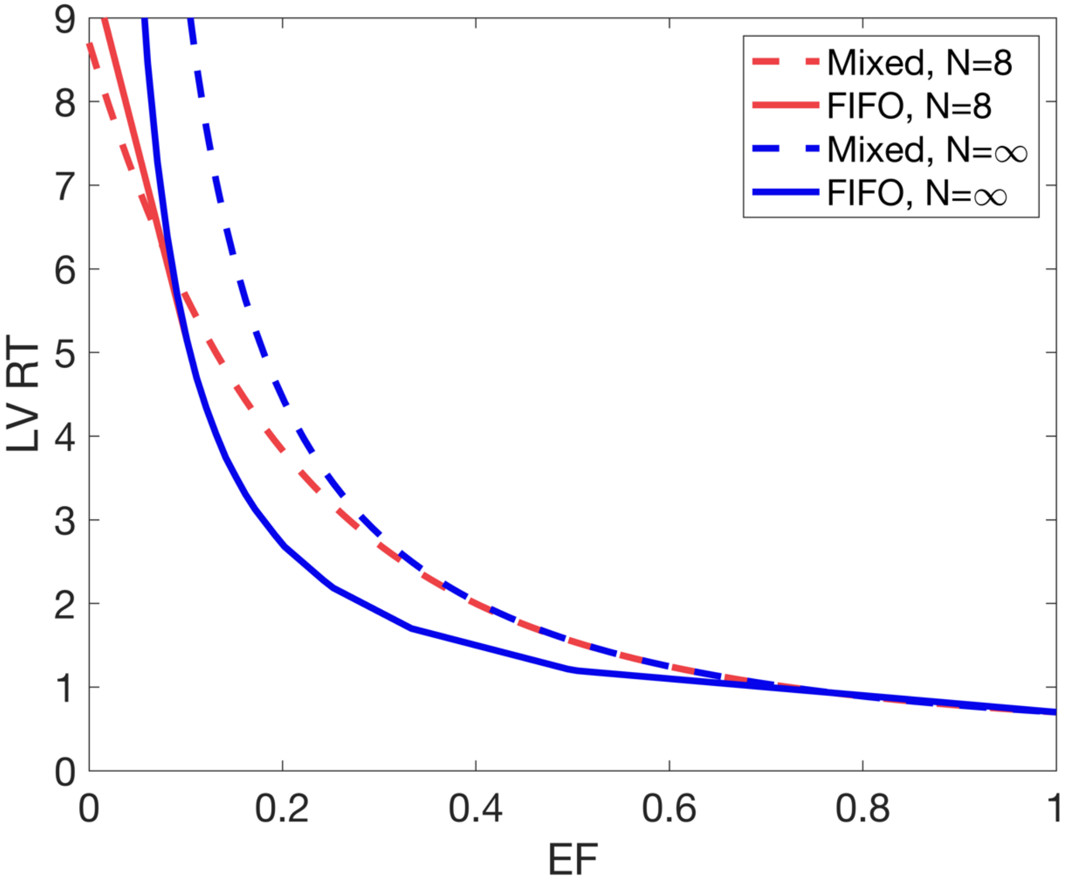


**Supplemental Figure 1:** Influence of the number of studied cycles in the calculation of LVRT using the perfect mixing model (dashed line) and the FIFO model (solid line) for N= 8 cycles (in red) and in the limit of infinity (in blue).
